# The hidden multiverse of ecological roles in Fungi

**DOI:** 10.64898/2026.07.14.738353

**Authors:** Carlos A. Aguilar-Trigueros, Sten Anslan, Amy Zanne, Nerea Abrego, Tessa Camenzind, Joseph D. Edwards, Jeff Powell, Adriana L. Romero-Olivares, Adam Frew

## Abstract

Ecological roles are treated as discrete, bounded units of biological organization, even as evidence shows that many taxa are versatile, performing more than one of them. Each record of versatility represents a boundary crossing, yet current categorical paradigms prevent this cumulative evidence from reshaping the roles themselves. We solve this by introducing the ecological multiverse, a self-correcting framework that turns boundary crossings into a network of role connectivity. Applied to Fungi, the multiverse recasts the major form of plant pathogenicity from an isolated disease role into a central hub linking much of fungal functional space, especially benign plant symbioses and decomposition. By exposing this topology, the multiverse transforms categorical views of ecological function into a dynamic map, revealing pathways of role change as organisms and knowledge evolve.

## Main Text

A central challenge in biology is organizing the complexity that emerges from the myriad interactions between living organisms, their environment, and one another (*1–3*). Ecology has traditionally approached this challenge by classifying organisms into discrete ecological roles based on how they acquire resources, occupy habitats, or influence the ecosystems in which they live (*4, 5*). These frameworks depict a mutually exclusive view of ecological organization and imply that eco-evolutionary processes constrain taxa to single roles (*6*). Yet nature is rarely so rigid. Across the tree of life, many organisms do not perform a single ecological role, but instead switch, combine, or perform different roles across space, time, and even life stages (*7, 8*). Each record of such versatility represents evidence that a predefined role boundary has been crossed. Nowhere is this more evident than in microbial systems, where short lifespans, rapid evolution, and abrupt shifts such as the emergence of pathogenicity from non-pathogenic lineages make ecological roles especially fluid. However, even as evidence of versatility accrues, we lack self-correcting systems to revise their own boundaries, allowing distorted boundaries in ecological organization to persist.
The challenge in revising these boundaries lies in uncovering the hidden connectivity structure among roles that emerges from ecological versatility. Without this topology, we cannot determine which roles have permeable boundaries, which represent isolated endpoints, or which pathways of role change are more plausible or constrained. To reveal this structure, we built the multiverse of ecological roles: a multidimensional landscape in which roles are linked by taxa capable of performing multiple roles. In this network, roles are nodes, versatile taxa create links, and accumulating evidence of versatility can strengthen, weaken, or redraw the boundaries among roles. Analogous to the diseasome in human genetics (*9*)—which exposed relationships among diseases through shared genes—we infer relationships among ecological roles through shared taxa. This approach allows functional structure to emerge from observed role sharing rather than rigid classification, providing a framework to quantify versatility, redefine ecological roles, and anticipate their change in a dynamic world.

### Revealing the hidden multiverse of ecological roles in the kingdom Fungi

The Kingdom Fungi provides an ideal system for building the first ecological multiverse because this group spans an extraordinary diversity of ecological roles often referred to as lifestyles (*10*), many of which shape planetary functioning (*11, 12*). These roles range from free-living decomposers that underpin nutrient and carbon cycling in soils (*13*) to pathogenic and mutualistic groups that regulate the fitness of myriad plant and animal hosts. Such interactions include mycorrhizal associations (the nearly universal mutualistic interaction between fungi and plant roots that enhance soil nutrient acquisition, promoting plant productivity (*14*)); the widespread association of fungal species with algae that forms lichens (*15, 16*); some of the most devastating plant pathogen diseases ever recorded (*17*); behavior-manipulating pathogens of insects (*18*), and rapidly evolving human diseases (*19*). These static labels expose the central problem of categorical classification: decades of evidence show that many fungal taxa switch, combine, or alternate among lifestyles (*20–22*), to the point that modern diversity surveys relying on eDNA reveals that most taxa found through next-generation sequencing (*23*) techniques belong to versatile taxa (Fig. 1). Yet this versatility remains largely recorded as annotation rather than used to redraw the boundaries among fungal roles. These boundaries are especially prone to distortion because they were not delineated through a systematic comparison of functional diversity, but emerged as historically contingent categories inherited from separate research traditions (*24*). For example, roles such as *arbuscular mycorrhiza* and *ectomycorrhiza* are defined within mycology, *plant pathogenicity* within plant pathology, and *human pathogen* within medical mycology (*25–28*). As a result, the same taxa can receive different ecological labels depending on disciplinary context. This patchwork of partially overlapping categories, each built on distinct criteria, has long hindered the synthesis of fungal functional data. Consequently, these static labels continue to propagate through databases, ecological models, and assessments of functional risk, including predictions of pathogenic roles in plants and animals (*17, 29–33*).

**Figure 1.**
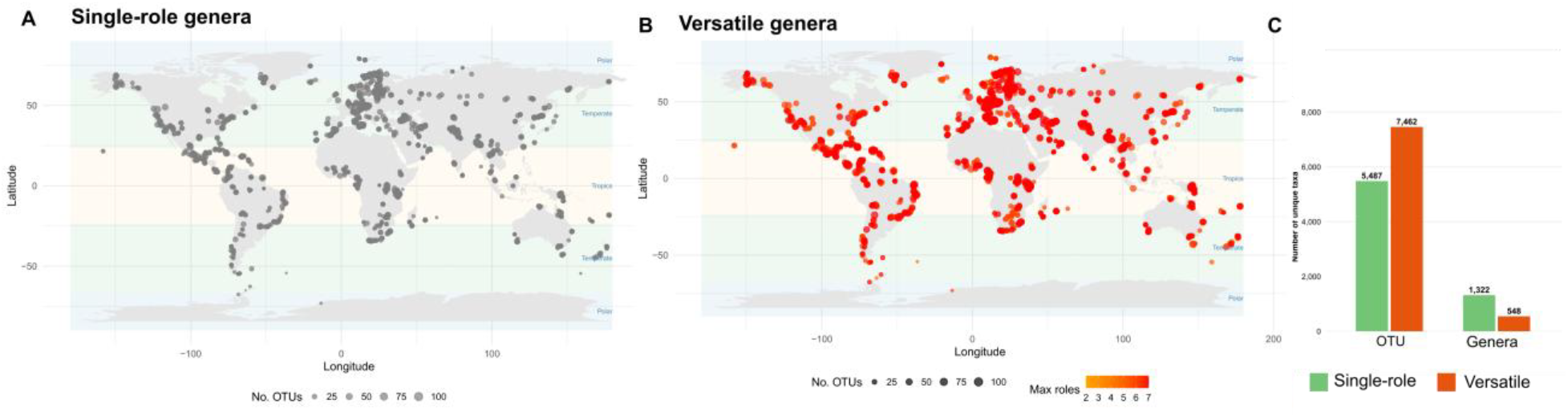
Occurrence and frequency of versatile taxa detected in eDNA surveys. Single-role taxa (a) and versatile taxa (b) exhibit broadly similar global occurrence patterns. Despite this overlap in distribution, most eDNA sequences are assigned to versatile taxa rather than single-role taxa.

To build this multiverse, we assembled 26 fungal roles by integrating databases reporting ecological roles (referred as lifestyles or guilds in this kingdom (Table S1)). We then use the topology of the ecological multiverse to determine where there is support for existing role boundaries and where it reveals they should be revised. If ecological versatility is rare, roles should remain isolated or form small, highly modular clusters, supporting the current view of ecological roles as discrete categories. If versatility is pervasive, roles should instead form a connected topology, indicating that current boundaries separate functions that taxa repeatedly bridge. A hub-dominated topology would further suggest that some roles are not isolated endpoints, but central phases through which many ecological roles connect; by contrast, a more uniform topology would indicate that versatility is broadly distributed across roles. Phylogenetic breadth adds a second layer for evaluating whether role boundaries hold. Weak clustering indicates that versatile taxa are distributed across the fungal phylogeny, showing that the same boundary is repeatedly crossed by distant lineages, providing stronger evidence that their current separation should be revised.

The fungal multiverse revealed a highly centralized topology in which ecological versatility was concentrated around a small set of plant-associated lifestyles (Fig. 2). This centralization produced very low modularity, with little separation among groups of roles (Fig. S1), indicating that fungal lifestyles do not organize into discrete, mutually exclusive categories. Necrotrophic plant pathogenicity, a disease strategy in which fungi kill host tissue to acquire nutrients (*34*)), emerged as the dominant hub, connecting to nearly all other roles. Rather than forming an isolated disease category, this role linked most frequently with plant endophytism, the asymptomatic colonization of living plant tissues, and with free-living saprotrophic lifestyles involved in the decomposition of dead organic matter. Nearly 50% of all versatile genera participated in necrotrophic plant pathogenicity, and roughly 35% linked this role with endophytism or saprotrophy. This concentration of versatility suggests that the boundary between plant-associated lifestyles and decomposer lifestyles is among the weakest in current fungal classification.

As these roles were also the most taxa-rich, it is possible that the prevalence of versatility simply reflects sampling effects (i.e. the more taxa, the more versatile taxa assigned). To account for this potential bias, we scaled the number of versatile genera against the total number of genera assigned to each ecological role (Fig. 2C). Accounting for this baseline relationship, versatility remained strongly concentrated around the same roles. Plant endophytism emerged as the strategy most dominated by versatile taxa (Fig. 2C, Fig. S1) (70% of genera and 30% of species are also associated with at least one additional strategy) followed by free-living saprotrophs and necrotrophic plant pathogenicity (with ∼30% of its ∼3,000 genera and ∼10% of its ∼10,000 species performing another ecological function)

**Figure 2.**
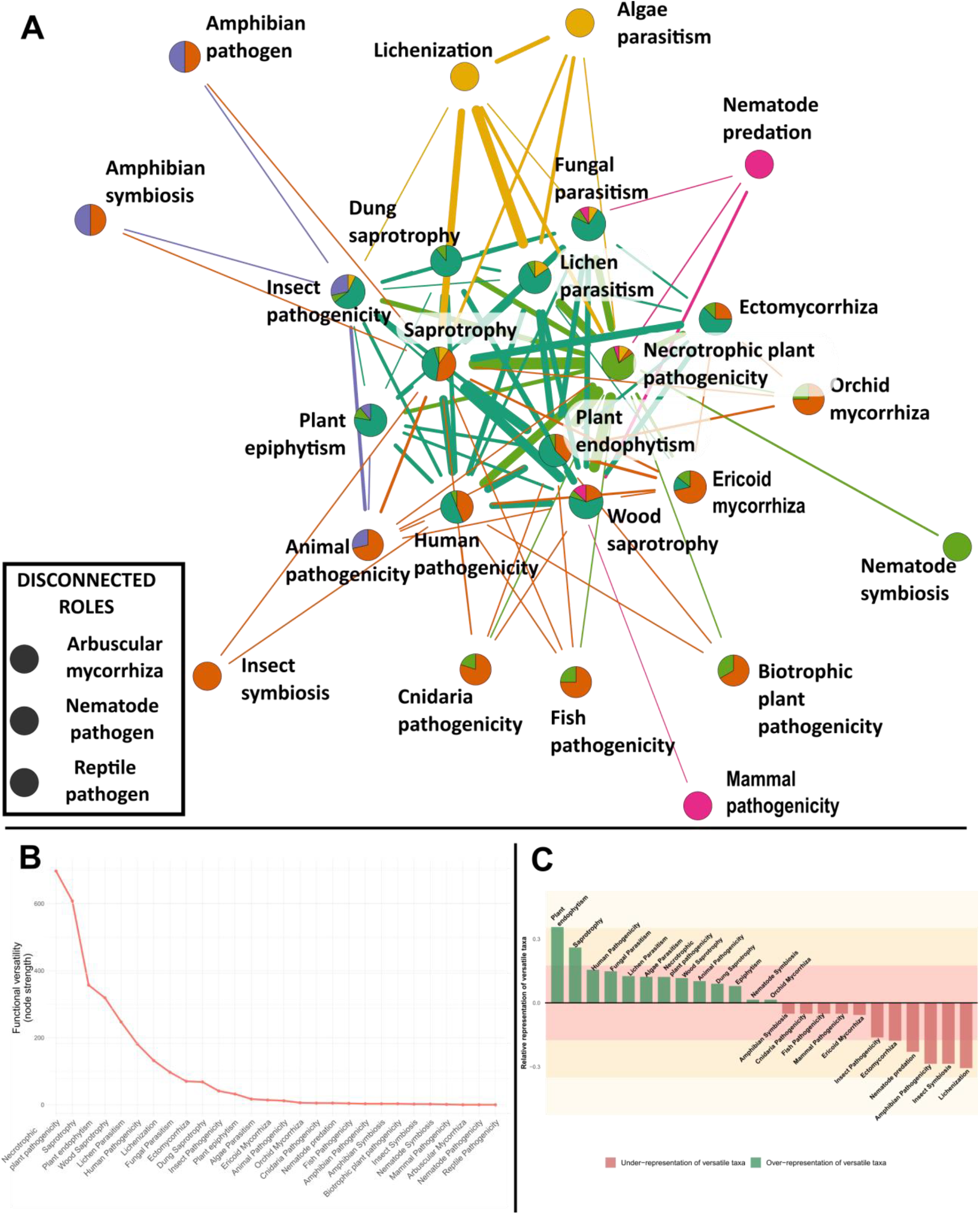
The fungal ecological multiverse. **(A)** Network of ecological roles connected through versatile genera. Pairs of roles are linked when at least two genera have been reported performing both. Line link thickness scales with the number of versatile genera connecting each pair while line link and pie colors indicate clusters of pairs identified with link similarity algorithms (see Fig. 3). The multiverse is highly connected and centralized, with necrotrophic plant pathogenicity, plant endophytism, and wood saprotrophy and saprotrophy acting as major hubs. **(B)** Magnitude of versatility across ecological roles, measured as node strength (the total number of versatile taxa associated with each role). **(C)** Ranking of ecological versatility after accounting for the scaling relationship between the number of versatile taxa and the total number of taxa assigned to each ecological role. Nodes with higher strength indicate the concentration of versatile taxa around that node in absolute values. The 0-line represents expected values based on the number of taxa assigned to a role; deviations from this line indicate the extent to which roles are dominated by versatile taxa (positive values) or specialists (negative values).

At the opposite extreme, some ecological roles showed less frequent cross boundaries. These roles emerged as weakly connected, or entirely disconnected nodes from the rest of the multiverse. The clearest case was arbuscular mycorrhiza, the mycorrhizal association formed by fungi in Glomeromycota (*14*)—which was completely disconnected from the rest of the network. Similarly, biotrophic plant pathogens (i.e., pathogens that acquire nutrients without inducing host tissue death (*35*)) showed extremely low versatility: among the 161 genera assigned to this role, only a single genus was associated with additional strategies (plant endophytism and human pathogenicity). Other symbiotic roles, including ectomycorrhizal fungi and lichenized fungi, were not fully isolated but remained weakly integrated into the multiverse. Lichenization was especially dominated by single-role taxa, with only 12% of its ∼905 genera and 1% of its ∼10,000 species associated with additional lifestyles. Animal-associated roles also showed evidence of bounded connectivity, although less strongly than symbioses with photosynthetic hosts. Insect pathogenicity, for example, was not isolated, but many of its connections clustered with other animal-associated symbioses, forming a distinct neighborhood reminiscent of the lichen-associated group. These patterns suggest that, unlike the more permeable boundary between plant pathogenicity, endophytism, and decomposition, some symbiotic roles represent more strongly bounded endpoints of fungal ecological organization.

### Ecological versatility reveals emergent functional neighborhoods

Because modularity among roles was low, we next asked whether the links themselves revealed recurrent patterns of boundary crossing. We analyzed the similarity structure among the 105 links of the ecological multiverse using a link-community approach (*36*). Unlike node-based clustering, which assigns each role to a single group, link communities group associations among roles, allowing the same role to participate in different functional neighborhoods depending on the roles with which it is connected. This provides a way to identify higher-order ecological organization without forcing roles into mutually exclusive categories.

This analysis revealed a primary division between links involving animal-associated roles and those involving plant-associated roles (Fig. 3). Insect endosymbiosis and nematode-trapping fungi formed some of the most distinctive association groups, indicating boundary crossings that are restricted to narrow regions of the multiverse. More broadly, animal-associated roles clustered with free-living saprotrophic and mycorrhizal roles, but less often with plant pathogenic or lichen-associated roles. Lichen-related roles, including lichenized fungi, lichen parasites, and algal parasites, formed a comparatively isolated neighborhood, with limited connections not only to animal-associated roles but also to other plant symbioses.

**Figure 3.**
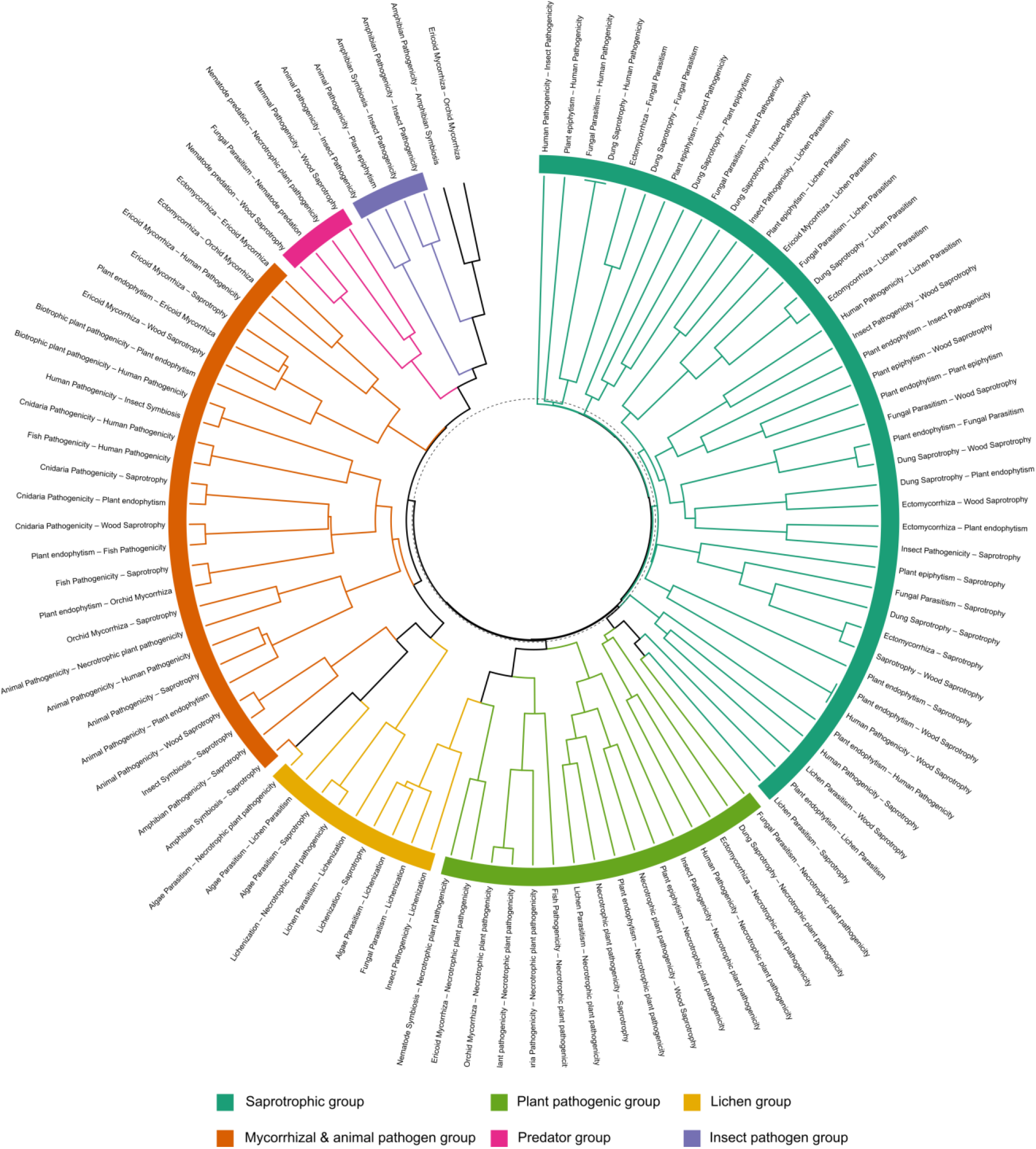
Hierarchical clustering of ecological links based on patterns of versatility. Each branch represents a network link, that is, a pair of ecological roles connected by versatile taxa. The six ecologicalgroups emerge based on the similarity of the neighborhoods of roles connected by versatile taxa (colored). The Insect group, the most distinct, connects insect pathogenicity with rare animal symbioses and contains no saprotrophic roles. The Predator group links nematode trapping with fungal parasitism and wood saprotrophy. The Mycorrhizal–animal pathogen group combines associations among mycorrhizal strategies and animal pathogens and is the most functionally diverse, encompassing 16 of the 26 roles. The Lichen group is dominated by links between lichenization and lichen parasitism. The two largest groups are the Plant pathogen group, comprising 16 roles linked primarily to the central hub of necrotrophic plant pathogenicity, and the Saprotrophic group, connecting general, wood, and dung saprotrophy with eight other roles.

Consistent with their high centrality, necrotrophic plant pathogenicity and saprotrophy formed two of the largest functional neighborhoods, linking to 17 and 11 roles, respectively. However, these neighborhoods differed in structure. The necrotrophic pathogenicity neighborhood was highly centralized, with many roles connected through this single hub but fewer connections among those roles themselves. In contrast, the saprotrophic neighborhood was more distributed, with links occurring more broadly among connected roles. Thus, link clustering turns accumulated evidence of ecological versatility into data-defined functional neighborhoods, rather than relying on predefined ecological categories.

### Phylogenetic clustering distinguishes broad boundary crossing from lineage-specific transitions

Phylogenetic structure provided a second layer for evaluating boundaries crossed by ecological versatility. Most links in the fungal multiverse showed some phylogenetic clustering, indicating that role sharing is often not randomly distributed across the fungal tree (Fig. 4). However, the strength of this clustering varied widely among links, revealing that some boundaries are crossed mainly by particular lineages, whereas others are crossed by distantly related taxa. Among the most central links involving necrotrophic plant pathogenicity, clustering was strongest for the connection with wood saprotrophy and weakest for the connection with plant endophytism. This contrast suggests that the boundary between necrotrophic plant pathogenicity and endophytism is broadly permeable across fungal lineages, whereas the connection to wood decomposition is more restricted, likely reflecting lineage-specific traits. A similar contrast emerged among lichen-associated roles: the frequent link between lichenization and lichen parasitism showed strong phylogenetic clustering, whereas the link between lichen parasitism and necrotrophic plant pathogenicity was much less clustered. Among links of intermediate centrality, the connection between ectomycorrhiza and saprotrophy showed especially strong phylogenetic clustering, indicating that this boundary is crossed by a narrower set of related taxa. Thus, even frequent boundary crossings can either reflect narrow lineage-specific transitions or broader connectivity across the fungal phylogeny.

**Figure 4.**
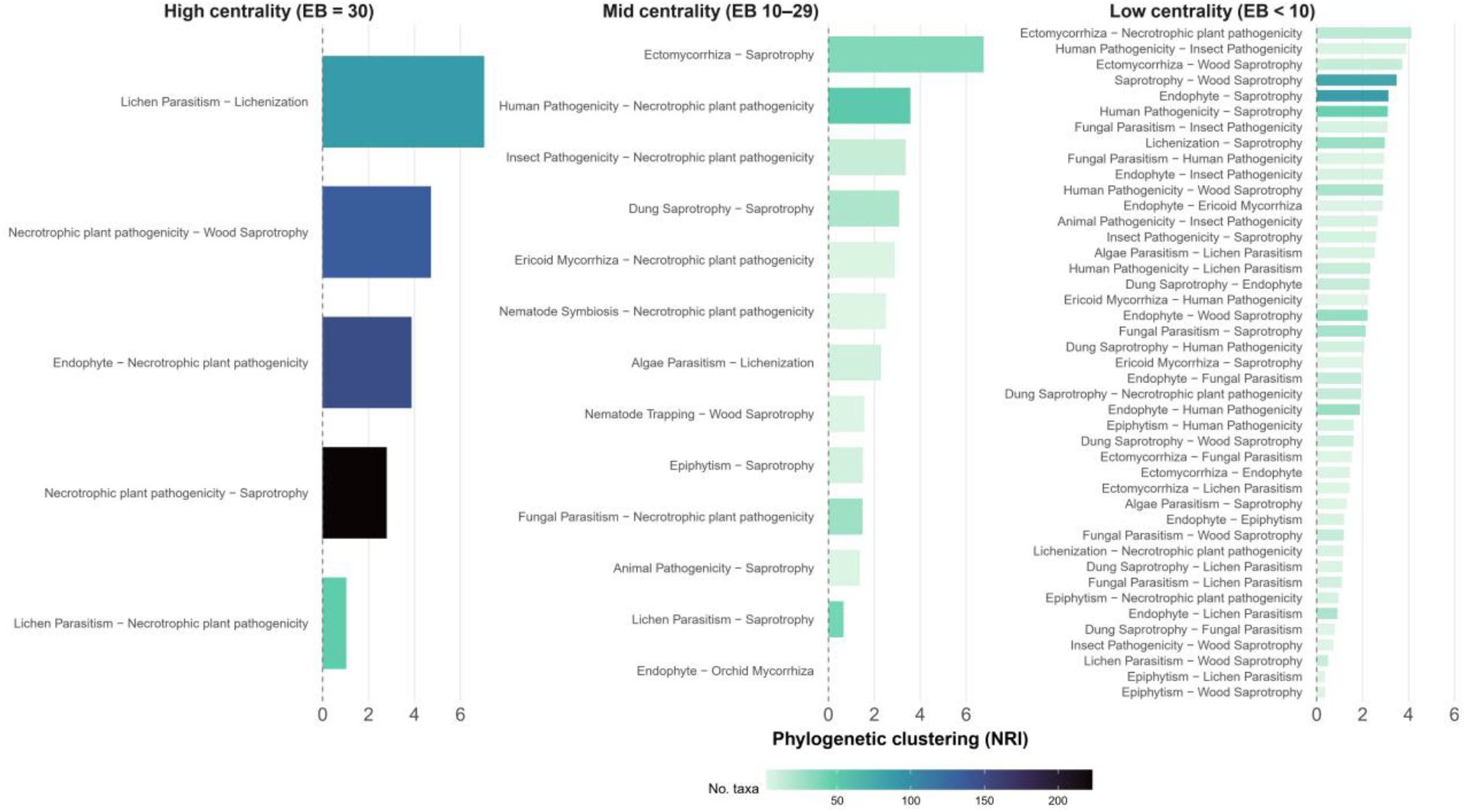
Phylogenetic clustering of links of the functional multiverse. Each link represents a versatile taxa (genera) that are reported to perform two ecological roles. Links with high-betweenness centrality form the backbone of the network containing the largest number of versatile taxa and connecting different clusters of roles. High phylogenetic clustering, measured with the net relatedness index, indicates that taxonomic diversity behind a link is concentrated in few clades out the whole fungal taxonomic tree. Given the mathematical relationship between edge betweenness centrality and NRI to the number of taxa, comparisons are most meaningful within each panel.

### Implications of the multiverse in redefining fungal ecological roles

The connectivity patterns revealed by the multiverse suggest that necrotrophic plant pathogenicity, saprotrophy, and plant endophytism are not discrete roles, but dynamic phases within a broader ecological strategy. This flexibility is likely underpinned by shared functional traits, particularly the use of cell wall–degrading enzymes (CWDEs), which enable fungi to exploit plant-derived substrates across living, stressed, senescent, and dead tissues (*37*). Because these enzymes underpin both saprotrophic and necrotrophic lifestyles, any taxa equipped with them can shift to either state as long as host defenses are overcome (in the case of the pathogenic role). Although CWDE activity can trigger host immune responses, and some necrotrophic pathogens have evolved mechanisms to suppress or evade them (*37, 38*), fungi can regulate their expression contextually (*39*). Their activation under host stress or senescence is therefore consistent with necrotrophic pathogen activity as a facultative or opportunistic state, rather than an isolated ecological role. The same logic may explain the link between endophytism and necrotrophy. Endophytes that grow asymptomatically within living tissues (*22*) can activate those enzymes as hosts senesce or become stressed, subsequently transitioning into saprotrophic phases (*21*). Reports of necrotrophic pathogens that persist as saprotrophs in the absence of hosts (e.g. overwintering) (*38*) further support the view of a broader strategy that allows fungi continuous resource use across their lifespan. Such strategy is congruent with the definition of versatility. The central position of necrotrophic pathogenicity in the multiverse is consistent with this intermediate phase linking resource use between living and dead substrates. Our findings are also consistent with fungal macroevolution patterns. Repeated transitions between saprotrophy and parasitism on photosynthetic hosts (*16*), together with evidence that enzymes for degrading autotroph-derived polymers were present near the origin of the kingdom (*40*), suggest that the weak boundaries among plant pathogenicity, endophytism, and decomposition reflect a long-standing feature of fungal ecology rather than a recent or isolated phenomenon. Resolving how widespread and mechanistically coordinated these strategies are across fungi remains an important direction for future work.

In contrast to highly connected roles, those with intermediate connectivity and stronger phylogenetic clustering represent partially bounded states. For example, the taxa rich link between lichen parasitism and lichenization, which captures a dynamic continuum between symbiosis and parasitism, remains phylogenetically clustered and confined to a restricted region of the multiverse.This pattern supports a lichen-associated functional neighborhood shaped by traits that evolved within particular fungal lineages (*41–43*). The transition between necrotrophic plant pathogenicity and wood saprotrophy also shows greater phylogenetic constraint and isolated position compared to transitions involving general saprotrophy (Fig. 4). Wood decomposition depends on specialized enzymatic machinery associated with lignin degradation, a relatively late evolutionary innovation, and is concentrated in a limited number of taxa (ref). Comparable patterns are observed with the mycorrhizal roles (except arbuscular ones), which form distinct, weakly connected neighborhoods with high phylogenetic clustering. The traits required for these interactions have also emerged in specific lineages and are associated with shifts in genetic makeup (*44*). Similar levels of specialization are documented in many insect-pathogenic fungi to infect their host, including, formation of haustoria in the infection process and the ability to manipulate host behavior to enhance spore dispersal, like in well described parasitic interactions of the fungal genus *Cordyceps* with insects (*18*). Together, these examples indicate that intermediate-connectivity roles represent better defined roles, in which the acquisition of novel traits expands ecological opportunity while simultaneously imposing trade-offs that reduce transitions to other roles.

At the other extreme of the connectivity gradient, roles with low or no connectivity—such as arbuscular mycorrhizal symbioses and biotrophic plant pathogenicity—behave as isolated endpoints with the best supported boundaries in the multiverse. Both biotrophic pathogens and arbuscular mycorrhizal fungi rely on complex cellular structures, such as haustoria and arbuscules, to extract nutrients from living host cells, and exhibit strict metabolic dependence on their hosts (*35*). These adaptations, which have evolved convergently across lineages, are also accompanied by reduced enzymatic capacity for independent resource acquisition (*45*) and loss to perform free living roles, and, as we show, any other role. Thus, these roles can be understood as relatively fixed states: specialized traits enable host dependence, but impose trade-offs that restrict transitions elsewhere in functional space. We also found several animal-associated symbioses that were completely disconnected from the multiverse, including reptile- and nematode-associated symbionts or pathogens. However, because each of these roles is currently represented by a single genus, it is premature to interpret their isolation as evidence of specialization. In contrast to well-characterized plant symbioses, animal-associated fungal symbioses remain poorly sampled, highlighting the need for more functional evidence to determine whether their apparent isolation reflects true specialization or limited knowledge.

### Revealing more ecological multiverses

We envision the ecological multiverse as a dynamic, self-correcting framework for advancing functional understanding across the tree of life. The fungal multiverse presented here is a first approximation, reflecting current knowledge based largely on expert-driven role annotations and incomplete taxonomic coverage. These limitations are unavoidable, but they are also central to the value of the framework: as more versatile taxa are reported and ecological annotations are revised, the connectivity of the multiverse can be recalculated, allowing classifications to change as evidence accumulates. We propose that, analogous to species hypotheses in DNA-based taxonomy(*46*), ecological roles can be treated as functional hypotheses: provisional units whose boundaries should be tested, supported, or redrawn as evidence accumulates. The current fungal multiverse already shows how roles grouped as plant pathogens can occupy opposite ends of the connectivity spectrum, from a dynamic phase embedded within broader ecological strategies and defined by weak boundaries, to an isolated ecological endpoint with well-supported boundaries. The multiverse therefore provides a systematic framework for revising ecological roles in light of the same evidence used to define them, in contrast to categorical systems that often remain static. We therefore encourage the continued expansion of functional annotations, particularly for microbial groups, and greater rigor and transparency in ecological classification, a frontier that still lags behind advances in taxonomy and phylogenetic systematics (*47*).

We further propose that the ecological multiverse is not a single map, but a family of maps operating across biological scales. The genus-level multiverse presented here is appropriate for fungi because genus is the resolution most commonly used to infer function in this kingdom, and because it captures role versatility accumulated over broad evolutionary timescales. Finer-resolution multiverses would reveal different dimensions of boundary crossing. Species- and strain-level multiverses could directly test within-organism role switching and environmental plasticity. Integrating these scales would allow short-term ecological versatility and long-term evolutionary change to be disentangled, turning the multiverse from a map of crossed boundaries into a framework for discovering the mechanisms that cross them.

The multiverse provides a framework for guiding more established -omic approaches. Although omic approaches reveal the traits that allow taxa to perform ecological roles (*48*), they are not designed to evaluate whether the functional categories being analyzed are the appropriate units of comparison. The multiverse adds this prior step by evaluating which role boundaries remain useful once ecological versatility is considered. This shift opens new questions, including why some traits promote versatility while others constrain it, and draws attention to understudied crossing—such as the link between wood saprotrophy and predatory behavior—whose mechanisms remain largely unexplored (*49*). In this way, the multiverse turns ecological roles into functional hypotheses that can be tested with increasingly mechanistic data. Finally, the multiverse shifts attention from ecological roles alone to the taxa that connect them. Versatile taxa not only challenge categorical systems; they represent a largely overlooked ecological phenomenon. By exploiting different resources across space and time, ecological versatility can buffer taxa against temporal or spatial resource limitation (*50*). As a consequence, these taxa may also become functional connectors in ecosystems: by performing multiple roles, they can influence several ecological processes and potentially contribute disproportionately to ecosystem dynamics, resilience, and the emergence of novel functions. Thus tracking versatile taxa through metabarcoding would make it possible to test whether ecosystems rich in such taxa are more resilient under environmental change.

Taken together, the ecological multiverse moves ecology from fixed categorical approaches toward a self-correcting view of functional organization. Categorical assignments—including functional types, trophic guilds, ecological strategies, functional groups, and lifestyles—have long provided essential structure for comparing organisms across the tree of life, from plants and animals (*4, 51–53*) to microbial diversity (*6, 54*). Yet when these categories remain fixed, they can preserve distorted boundaries even as evidence accumulates that taxa repeatedly cross them. This limitation is especially acute for microbial diversity, where functional categories are often inferred from incomplete annotations and inherited from classification schemes developed for other groups. Given the central role of microbes in regulating ecosystem processes, improving how microbial functions are defined, tested, and revised is essential for advancing ecological prediction. The multiverse resolves this problem by treating ecological roles as functional hypotheses: provisional units whose boundaries can be tested, supported, weakened, or redrawn. By converting overlap among roles into a map of functional space, the multiverse reveals how ecological functions are connected, which boundaries remain meaningful, and which pathways of role change may shape the organization and evolution of life as organisms, and our knowledge of them, evolve.

## Acknowledgments

We thank comments by Dirk Brockmann on link community analysis. We thank Isin Kosemen for developing interactive tool for this project.

## Author contributions

Conceptualization: CAA-T

Methodology: CAA-T, AF, SA

Investigation: CAA-T, AF

Visualization: CAA-T

Funding acquisition: CAA-T

Supervision: CAA-T, AF

Writing – original draft: CAA-T, AF, SA, AZ, JP, TC

Writing – review & editing: CAA-T, AF, SA, AZ, JP, JDF, NA, AR-O, TC

## Data, code, and materials availability

All data, code, and materials used in the analysis will be made available in open source repositories

