## Supplementary material for "The hidden multiverse of ecological roles in Fungi": Materials and methods

**The PDF file includes:**

Materials and Methods

### Materials and Methods

#### Data acquisition

To map connectivity among fungal ecological roles, we integrated information from sources spanning mycological systematics, plant pathology, medical mycology, and ecological subfields. We first assembled species-level role annotations from FunGuild (*1*), which remains a widely used reference for functional annotation of fungi in ecological community surveys. To expand the taxonomic and functional coverage beyond FunGuild, we incorporated annotations from specialized resources: the USDA host-fungus database (https://fungi.ars.usda.gov/), accessed with the R package `rusda` (*2*), for plant-pathogenic fungi; LIAS for lichenized and lichenicolous fungi (http://liaslight.lias.net/) ; DEEMY for ectomycorrhizal fungi (http://www.deemy.de/); TraitAM for arbuscular mycorrhizal fungi (*3*); and published compilations for entomopathogenic and human-pathogenic fungi (*4*, *5*). We standardized to a common taxonomy across all thse sources following accepted names in Catalogue of Life (*6*) while we standardized ecological role names using FunGuild nomenclature as the reference. We further divided the plant pathogen category into biotrophic and necrotrophic groups. Biotrophic pathogens are obligate symbionts that require living host tissue to complete their life cycle, whereas necrotrophic pathogens induce host tissue death to acquire nutrients and complete infection (*7*, *8*). When applicable, we also annotated the specific disease types associated with plant-pathogenic taxa. Following the USDA host-fungus database and ref. (*9*), for biotrophic plant pathogens those diseases included “rusts”, “smuts”, “powdery mildews”, and “black mildews”, whereas for necrotrophic pathogens diseases included “cankers”, “leaf spots”, “scorch”, “anthracnose”, “blotch”, “blight”, “damping-off”, “rots”, and “necrosis”.

#### Assembling the ecological metaverse

Species-level ecological assignments were collapsed into a genus-by-role matrix, with genera as rows and ecological roles as columns. For each genus, we aggregated all ecological roles assigned to any species within that genus. Thus, if one species in a genus was annotated as a plant pathogen and another as a saprotroph, the genus received both role assignments and was classified as versatile. Matrix entries were binary, indicating whether at least one species within a genus was assigned to a given role. We used genus-level aggregation because genus is the most common taxonomic resolution used to infer fungal function in ecological studies. In this framework, versatility reflects the extent to which ecological role boundaries are crossed within closely related taxa.

We transformed the genus-by-role matrix into a role-by-role adjacency matrix, with ecological roles represented as both rows and columns. The diagonal of this matrix indicates the number of genera assigned to each role, whereas off-diagonal elements indicate the number of genera assigned to both roles. We used this matrix to construct a weighted graph in which nodes represent ecological roles, edges indicate that at least one genus links two roles, and edge weights correspond to the number of genera shared between those roles. In the visual representation of the network, thicker edges indicate stronger connections, i.e., a larger number of genera assigned to both roles. This framework also allows the identification of versatile genera, defined here as genera assigned to more than one ecological role. These genera do not form nodes in the role network; rather, they define the links among roles and therefore determine the connectivity structure of the ecological multiverse

Identifying roles most frequently crossed by ecological versatility

For each ecological role in the network, we calculated node degree and node strength to quantify how permeable its boundaries were to ecological versatility. Node degree was defined as the number of other roles to which a focal role was connected through versatile genera. Thus, roles with high degree have boundaries that are crossed toward many different parts of functional space. Node strength was defined as the sum of edge weights connected to a focal role, and therefore reflects the total number of versatile genera linking that role to others. Roles with high node strength have boundaries crossed by many genera, even if those crossings involve only a limited number of other roles.

Because larger roles are expected to contain more versatile genera simply by chance, we also tested whether some roles contained disproportionately more or fewer versatile genera than expected from the number of annotations to that role. That is, for each role, we quantified role size as the total number of genera assigned to that role and versatility as the number of genera assigned to that role and at least one additional role. We then fitted a power-law relationship between role size and the number of versatile genera. This scaling relationship provided a baseline expectation for how versatility increases with role size. Roles with positive deviations from this relationship contained more versatile genera than expected, indicating more permeable boundaries after accounting for role size, whereas roles with negative deviations contained fewer versatile genera than expected, indicating more strongly bounded roles. The scaling relationship and its significance were tested using standardized major axis regression implemented with the `sma` function in the `smatr` R package (Warton et al., 2012)

Global distribution of versatile taxa

To assess the global occurrence of ecologically versatile fungi, we integrated our lifestyle database with the Global Soil Mycobiome Consortium (GSMc) dataset, which comprises 722,682 OTUs sampled across 3,068 soil samples worldwide. Each OTU was first matched to genus level using the GSMc taxonomy file, and only genus-annotated OTUs were retained (16,359 OTUs). Ecological role information was then assigned to each genus by joining the GSMc taxonomy against our compiled ecrological role database. Genera classified to a single ecological role were designated as *single-role* taxa, while genera with evidence of two or more roles were designated as *versatile* (ecologically multifunctional) taxa.

Definition of higher order functional regions in the multiverse

To identify higher-order structure in the ecological multiverse, we used two complementary approaches. First, we applied the fast greedy modularity optimization algorithm (*10*) implemented in the R package igraph (*11*) to determine whether boundary crossings among ecological roles were organized into discrete modules (i.e. groups of ecological roles that were more densely connected to one another than to the rest of the network). This node-based approach identifies groups of roles that are more densely connected to one another than to the rest of the network. In the context of the multiverse, such modules would indicate that, although ecological versatility crosses categorical boundaries, these crossings remain confined to a narrow set of roles, forming relatively closed units that could be interpreted as higher-order ecological roles.

Because modularity was low, we then used a link community approach to identify more flexible forms of organization in the multiverse. Rather than assigning each ecological role to a single module, link community analysis clusters the connections among roles. This approach is useful when the same role participates in multiple regions of functional space, because roles can inherit multiple memberships through their different links. We therefore used link similarity analysis (*12*) implemented in the R package linkcomm (*13*) to identify functional neighborhoods: groups of role-to-role links that share similar local network context, even when the roles themselves do not form discrete modules. In this framework, functional neighborhoods emerge from recurrent patterns of boundary crossing. For this analysis, functional role pairs shared by a single genus were excluded from the link-community analysis, as single-taxon links form trivial singleton communities and do not contribute to the partition density maximisation used by the algorithm (Ahn *et al.* 2010; Kalinka & Tomancak 2011).

Estimating the phylogenetic breadth of boundary crossings

To assess whether crossings between ecological role boundaries were phylogenetically restricted or broadly distributed across fungi, we estimated the phylogenetic structure of the genera supporting each role-to-role link. We first constructed a taxonomy-based phylogenetic tree including all genera in the ecological multiverse. Although taxonomy-based trees lack precise branch-length information, they provide a broad approximation of evolutionary relationships and are useful for evaluating large-scale phylogenetic structure across taxonomically diverse datasets (*14*).

For each link in the multiverse, we identified the set of versatile genera assigned to both ecological roles connected by that link. We then tested whether these genera were more phylogenetically clustered or more dispersed than expected under a null distribution using the net relatedness index (NRI). In this context, strong phylogenetic clustering indicates that a boundary is crossed mainly by closely related genera, suggesting that the connection between roles may depend on lineage-specific traits. Weak clustering indicates that the same boundary is crossed by distantly related genera, providing stronger evidence that the separation between those roles is broadly permeable across the fungal phylogeny.

The taxonomy-based tree was constructed using the `as.phylo` function in the `ape` package (*15*). Phylogenetic structure was estimated using the `ses.mpd` function in the `picante` package (*16*), which compares the observed mean pairwise phylogenetic distance among genera supporting each link to a null distribution

1. N. H. Nguyen, Z. Song, S. T. Bates, S. Branco, L. Tedersoo, J. Menke, J. S. Schilling, P. G. Kennedy, FUNGuild: An open annotation tool for parsing fungal community datasets by ecological guild. *Fungal Ecology* **20**, 241–248 (2016).

2. F.-S. Krah, C. Bässler, C. Heibl, J. Soghigian, H. Schaefer, D. S. Hibbett, Evolutionary dynamics of host specialization in wood-decay fungi. *BMC Evol Biol* **18**, 119 (2018).

3. V. B. Chaudhary, L. F. Nokes, J. B. González, P. O. Cooper, A. M. Katula, E. C. Mares, S. Pehim Limbu, J. N. Robinson, C. A. Aguilar-Trigueros, TraitAM, a global spore trait database for arbuscular mycorrhizal fungi. *Sci Data* **12**, 588 (2025).

4. J. P. M. Araújo, D. P. Hughes, "Diversity of Entomopathogenic Fungi" in *Advances in Genetics* (Elsevier, 2016; https://linkinghub.elsevier.com/retrieve/pii/S0065266016300013)vol. 94, pp. 1–39.

5. L. Irinyi, C. Serena, D. Garcia-Hermoso, M. Arabatzis, M. Desnos-Ollivier, D. Vu, G. Cardinali, I. Arthur, A.-C. Normand, A. Giraldo, K. C. da Cunha, M. Sandoval-Denis, M. Hendrickx, A. S. Nishikaku, A. S. de Azevedo Melo, K. B. Merseguel, A. Khan, J. A. Parente Rocha, P. Sampaio, M. R. da Silva Briones, R. C. e Ferreira, M. de Medeiros Muniz, L. R. Castañón-Olivares, D. Estrada-Barcenas, C. Cassagne, C. Mary, S. Y. Duan, F. Kong, A. Y. Sun, X. Zeng, Z. Zhao, N. Gantois, F. Botterel, B. Robbertse, C. Schoch, W. Gams, D. Ellis, C. Halliday, S. Chen, T. C. Sorrell, R. Piarroux, A. L. Colombo, C. Pais, S. De Hoog, R. M. Zancopé-Oliveira, M. L. Taylor, C. Toriello, C. M. De Almeida Soares, L. Delhaes, D. Stubbe, F. Dromer, S. Ranque, J. Guarro, J. F. Cano-Lira, V. Robert, A. Velegraki, W. Meyer, International Society of Human and Animal Mycology (ISHAM)-ITS reference DNA barcoding database—the quality controlled standard tool for routine identification of human and animal pathogenic fungi. *Medical Mycology* **53**, 313–337 (2015).

6. O. Bánki, Y. Roskov, M. Döring, G. Ower, D. R. Hernández Robles, C. A. Plata Corredor, T. Stjernegaard Jeppesen, A. Örn, T. Pape, D. Hobern, S. Garnett, H. Little, R. E. DeWalt, J. Miller, T. Orrell, R. Aalbu, J. Abbott, C. Abreu, A. Acero P, O. Acevedo-Charry, T. Adriaens, C. Aedo, E. Aescht, B. Agboola, D. Agosti, H. D. Agudelo-Zamora, W. Ahlmer, S. Ahyong, M. Akhoundi, C. Aldea, D. Alderman, S. Alexander, M. A. Alonso-Zarazaga, B. Alvarez, N. Ananjeva, P. Anastasiu, R. Anderson, A. J. Andrade, G. C. Andrella, M. Anions, Anonymus, D. Z. Antić, L. S. Antonietto, C. Arango, M. Arianoutsou, J. A. Arias-Buriticá, E. Arriaga-Varela, T. Artois, O. Ascuntar-Osnas, M. Atahuachi Burgos, S. Atkinson, J. J. Atwood, S. Augustin, J. E. Avendaño, J. E. Avendaño C., S. Bacher, M. Backlund, Â. L. Bagnatori Sartori, N. Bailly, J. Baixeras, E. Baker, A. Balan, E. Balletto, R. Bamber, S. Bandyopadhyay, A. Barber, H. Barber-James, R. Barbosa Pinto, R. Barrett, M. Barros-Barrios, L. Bartolozzi, I. Bartsch, A. Baya, C. Başnou, G. Beccaloni, J.-N. Beisel, C. L. Bellamy, D. Bellan-Santini, P. F. Bellinger, B. S. Beltrán León, Y. Ben-Dov, L. Benichou, H. Berlanga, M. F. Bermúdez-Higinio, J. Bernot, S. Bertolino, J. Beta, G. Biffi, I. Blasco-Costa, J. S. Boatwright, P. Bock, J. D. Bogota-Gregory, H. Bohn, B. Bolton, L. Bonesi, L. M. Borges, P. Borges, R. Bortoluzzi, R. L. Bossard, C. Bota-Sierra, J. P. Botero, P. Bouchard, N. Boury-Esnault, Y. Bousquet, R. Bouzan, G. Boxshall, C. Boyko, S. Brandão, H. Braun, R. Bray, G. Brazil Flora, G. Brehm, F. Bretagnolle, J. C. Brinda, P. D. Brock, S. L. Broich, D. Brosens, L. Brouillet, J. Brown, S. Brown, S. Brullo, A. Bruneau, L. Bush, T. Büscher, M. Błażewicz-Paszkowycz, A. Cabras, E. Cacabelos, S. Cairns, I. Calabuig, R. Calderón-Parra, M. Calonje, I. S. Campos-Filho, W. Cardinal-McTeague, J. Cardona-Duque, D. Cardoso, L. Cardoso, A. B. G. de L. Carvalho, P. Carvalho, A. Castaño

Rivera, R. C. Castilho, E. B. Castro, I. C. Castro Silva, A. Cervantes, W. R. Chamorro-Fuertes, J. L. Chapuis, B. Chauvel, A. Chernyshev, H. Chevillotte, F. Chiron, L. M. Choo, H. Choong, K. A. Christiansen, F. Cianferoni, P. Cifuentes-Ruíz, M. M. Cigliano, C. X. Cisteil Xinum, R. Clarke, J. Clavijo-Bustos, P. Clergeau, T. Cobra e Monteiro, A. Collins, S. Collins, J. Compton, J. Cooper, D. Copilaş-Ciocianu, L. Corbari, R. Cordeiro, CoreoideaSF Team, K. Cortés-Hernández, M. J. Costello, F. Coursol, S. Crameri, J. Creuwels, J. A. Cruz-López, A. M. Cuervo, T. Cunha, P. Cárdenas, B. Dagley, M. Daly, M. Daneliya, R. Dangi, J.-C. Dauvin, P. Davie, A. Davies, C. De Broyer, S. De Grave, H. C. De Lima, J. De Prins, W. De Prins, F. De Sousa, M. De la Estrella, R. DeSalle, P. Decker, W. Decock, A. Delgado-Salinas, P. Delipetrou, C. Deliry, P. M. Dellapé, J. Den Heyer, P. Desmet, M.-L. Desprez-Loustau, S. Devin, F. Di Giovanni, V. Didžiulis, W. Diewald, K.-D. Dijkstra, V. Dincă, D. A. Dmitriev, C. DoNascimiento, M. Dohrmann, D. Donsker, Ó. Dorado, F. Dorkeld, R. Downey, L. Duan, R. Duno de Stefano, F. Dämmrich, J. Dépaquit, M. Détaint, M.-C. Díaz, D. C. Eades, M. Ebert, M. Á. Echeverri-Galvis, A. N. Egan, M. Eitel, A. El Nagar, M. Eleaume, C. C. Emig, M. S. Engel, H. Enghoff, P. Esquete Garrote, F. Essl, F. A. Estela, G. A. Evans, N. L. Evenhuis, M. Falcão, Z. Faltynek Fric, F. Farruggia, K. Fauchald, D. Fautin, M. Favreau, C. Favret, D. Fenner, J. Ferm, B. Figuerola, FinBIF, B. Fisher, C. Fišer, A. Fonseca-Cortés, S. Fontinha, L. Forró, A. P. Fortuna-Perez, H. Fortune-Hopkins, A. Franquinho Aguiar, P. Fritsch, A. Fuchs, S. Fujimoto, H. Furuya, E. Gagnon, E. Galati, H. Galea, B. S. Galil, K. García, L. García-Prieto, O. Gargominy, R. Garic, R. Gasca, H. J. Gasca Álvarez, J.-L. Gattolliat, M. Geiger, M. Geiser, P. Genovesi, G. Georgiev, T. Georgiev, S. Gerken, F. Gherardi, A. Gibau de Lima, D. Gibson, C. Gielis, F. Gill, T. Gilligan, G. Giribet, J. C. Girón Duque, A. Gittenberger, G. Giusso del Galdo, S. Gofas, A. Golikov, S. Gollasch, C. Gomez, M. Goncharov, A. I. Gondim, A. González-Alvarado, M. González-Córdoba, C. Goodwin, R. Govaerts, M. Grabowski, A. de A. Granado, B. de S. Gregório, J. R. Grehan, R. Grether, D. A. Grimaldi, O. Gross, J. M. Guerra-García, A. Guglielmone, E. Guilbert, L. Guillou, T. Guldberg Frøslev, W. Gurumse, J. Gusenleitner, G. Guzmán, C. Gyeltshen, C. Gómez, H. Gómez de Silva, M. Göker, Y. HAN, F. Haas, M. Haas, K. A. Hadfield, E. Hajdu, M. Hassler, M. W. Hastriter, A. Hausmann, B. W. Hayward, M. Hejda, L. Hendrich, E. Hendrycks, D. Hennen, T. J. Henry, F. A. Hernandes, D. Hernandez, F. Hernandez, D. Hernández, H. Hernández Macías, J. C. Hernández-Crespo, E. E. Herrera-Collazos, G. Herrera-R, D. Hincapié-Montoya, D. M. Hincapié-Montoya, A. Hine, B.-C. Ho, A. Hodson, B. Hoeksema, M. Hoenemann, J. Holstein, M. Hooge, J. Hooper, H. Hopkins, M. Hopkins, I. Horak, T. Horton, J. Hošek, C. Hughes, L. Hughes, T. Hughes, P. E. Hulme, R. Huys, J. Háva, C. Häuser, H. Höfer, M. Hüne, J. Iganci, L. F. M. Iniesta, Institute of Zoology Chinese Academy of Sciences, International Union for Conservation of Nature, T. Ishikawa, J. Ishong, L. JI, C. Jackson, F. Janssens, R. Jardim, R. Jaskuła, D. Jaume, F. Javadi, K. Jazdzewski, J. Jenkins Shaw, C. D. Jersabek, K. P. Johnson, M. A. Johnston, L. Jordão, M. Josefsson, P. Jóźwiak, H. Kajihara, K. Kakui, A. Kallies, M. J. Kamiński, K. Kanda, S. Kark, J. Kathirithamby, S. Katisho, K. Kauhala, M. Kelly, M. Kenis, R. Khalikov, Y.-H. Kim, R. King, L. Kirao, P. Kirk, I. Kitching, M. Klautau, N. Klazenga, R. Kleukers, B. B. Klitgaard, S. Klotz, N. Kmentova, M. Kobelt, S. Koenemann, N. M. Korovchinsky, R. Kostova, A. Kotov, T. Kramina, T. Krapp-Schickel, A. Kremenetskaia, K. Krishna, V. Krishna, A. Kroh, A. S. Kroupa, A. Kury, A. B. Kury, M. S. Kury, J. Kvaček, I. Kühn, C. LIN, B. LIU, O. Lachenaud, A. Ladino-Peñuela, C. Lado, G. Lamas, P. W. Lambdon, G. Lambert, L. Lana C. Atunes, T.-B.

Larsson, M. Lavin, D. Lazarus, F. Le Coze, M. Le Roux, S. LeCroy, J. Ledis Linares, H. Lee, S. Lee, M. F. Leitner, N. G. Leuro Robles, G. P. Lewis, S.-J. Li, Y.-H. Li, J. Li-Qiang, R. (†) Lichtwardt, S.-C. Lim, D. Lindsay, H. Liu, V. Lohrmann, S. J. Longhorn, C. Lopez-Vaamonde, W. Lorenz, O. Lorvelec, J. Lowry, M. Loyer, F. Lozano, V. Lukhtanov, R. Lumen, C. H. Lyal, P. López-Bedoya, A.-N. Lörz, K. MA, D. Maes, P. Magnien, C. Mah, I. Makhov, N. Mal, V. Malécot, T. Mamos, R. Manconi, V. Mansano, H. Marchante, K. Markello, K. Martens, J. H. Martin, P. Martin, C. A. Martínez Muñoz, D. E. Martínez-Revelo, K. S. Mashego, S. Maslakova, B. Maslin, S. Mattapha, M. Mańko, C. McFadden, S. McKamey, J. A. McMurtry, S. Meades, J. Means, C. A. Medina-Uribe, M. A. Medrano, J. Mees, J. P. Meier-Kolthoff, K. Meland, I. Melo, A. C. Mendes, B. Mendoza-Garfias, M. Menezes de Sequeira, K. Merrin, N. C. Mesa, C. Messing, C. G. C. Mielke, A. Migeon, L. J. Migliore, D. R. Miller, C. Mills, K. Milto, D. Minchin, A. Minelli, L. S. Miranda, V. Mironov, T. Mita, D. Mitchell, MolluscaBase eds., T. Molodtsova, J. C. Monje, J. F. Montenegro Valls, A. Montgomery, R. Mooi, D. M. Morales-Martínez, A. Morandini, R. Moreira da Rocha, I. Moreu, C. Morrow, A. Moteetee, M. L. Munguira, L. Murillo-Ramos, B. Murphy, T. Mwadime, J. P. Z. Narita, E. A. Nascimento, M. Nebel, K. Neill, J. C. Neita Moreno, W. Nelson, Nemys eds., W. Nentwig, A. I. Neto, U. Neu-Becker, T. A. Neubauer, B. Neuhaus, A. Newton, P. Ng Kee Lin, A. Nguyen, D. Nicolson, J. E. Nielsen, A. Nijhof, T. Nishikawa, M. A. Niño-Suárez, J. Norenburg, L. Novoa, S. A. O'Donnell, P. O'Grady, T. O'Hara, D. Obondo, A. Occhipinti-Ambrogi, J. Ochieng, R. Ochoa, H. Ohashi, K. Ohashi, E. Okposio, S. Olenin, I. Olenina, A. Oliveira, P. Oliveira, J. Ollerenshaw, P. Oosterbroek, D. Opresko, A. Ortega-Lara, R. Ortega-Álvarez, R. Osborne, H.-J. Osigus, J. D. Oswald, Y. Ota, D. Otte, D. Ouvrard, I. Ovcharenko, M. PIAO, L. Paganucci de Queiroz, J. S. Palacios Rodríguez, A. Pandey, V. E. Panov, M. I. Parente, A. C. Parte, M. Pascal, G. Paulay, D. Paulson, L. Penev, R. T. Pennington, J. da S. Pereira, S. G. G. Pereira, D. Perez-Gelabert, J. Pergl, I. Perglová, F. Perveen, A. Petrusek, P. Phillipson, D. Pica, U. Pinheiro, J. Pino, M. Pires Morim, A. Pisera, B. Pitkin, C. Plata, D. Plotkin, B. Poatskievick Pierezan, A. Polanco, A. Polanco F., G. Poore, M. Povydysh, K. Prasad, R. A. Praxedes, B. Price, J. Prudhomme, W. J. Pulawski, R. Pyle, P. Pyšek, C. X. Pérez-Hernández, F. Pühringer, W. Rabitsch, H. Rajaei, N. Rakotonirina, G. Ramos, H. E. Ramírez-Chaves, J. Rando, F. J. Randrianambinintsoa, F. Ranzato Filardi, C. Raper, P. Rasmussen, J.-Y. Rasplus, S. Ratcliffe, B. Rathod, L. Raz, G. Read, D. M. Reeder, T. Rees, M. Reich, J. D. Reimer, L. C. Reimer, J. O. Rein, A. Rendón Ramírez, J. Reynolds, L. Reyserhove, J. Rincón, M. Rius, T. Robertson, G. Robinson, G. S. (†) Robinson, E. Rodríguez, W. D. Rodríguez, V. Rodríguez-Contreras, A. Roques, S. Rosenfeld, D. Roy, H. Roy, M. Ruggiero, P. Ríos, Z. SHA, L. M. Sabater, R. Sabroux, J. Salmela, K. Samimi-Namin, A. Sanborn, M. K. Sandoval-Espinel, M. Sanjappa, A. Santos-Guerra, J. Sardà Carbasse, M. Sartori, K. Sattler, G. Sauter, D. Sauvard, R. Scalera, B. Schierwater, S. Schilling, R. Schley, C. Schmid-Egger, S. Schmidt, A. Schmidt-Rhaesa, P. Schoolmeesters, M. Schorr, B. Schrire, P. Schuchert, R. T. Schuh, C. Schönberg, R. Schütz Rodrigues, M. Scoble, G. Seijo, E. P. Seleme, A. Senna, C. Serejo, A. Serrano, A. Sforzi, K.-T. Shao, N. Shenkar, T. A. Shiganova, P. Shimabukuro, G. Shimizu, S. Shirley, A. Shwartz, V. Siegel, P. Sierwald, P. Sihvonen, D. Sikes, A. Silva Flores, C. Silva de Carvalho, M. Sim-Sim, E. Simmons, M. F. Simon, T. Simonsen, C. E. Simpson, N. Simões, S. Sinev, F. Sinniger, Y. Sirichamorn, Y. P. Sissa-Dueñas, L. Skipper, M. Skvarla, I. Smirnov, R. Smirnov, A. Smith, A. D. Smith, V. S. Smith, D. Soares Gissi, D. Sokoloff, W. Solarz, S. Sotuyo, J. Souma, E. J.

South, D. Souza, J. F. Souza-Filho, L. Spearman, J. Spelda, S. Stefanelli, A. Steiner, W. Steiner, T. Stemme, W. Sterrer, D. Stevenson, M. B. D. Stiewe, F. G. Stiles, C. H. Stirton, S. Straub, G. Stueber, S. Stöhr, S. Subramaniam, B. Swalla, J. Swedo, H. A. B. Sá, S. Sáfián, N. Sánchez Santos, L. A. Sánchez-González, M. Sánchez-Ruiz, C. Sérgio, M. V. Sørensen, C. A. Taboada-Verona, D. M. Takiya, A. H. Tandberg, G. Tavakilian, J. J. Tavera Vargas, K. Taylor, The Danish Agency for Green Land Use and Aquatic Environment, A. Thessen, D. Thomas, J. D. Thomas, P. Thomas, S. Thomson, S. Thorpe, ThripsWiki, E. Thuesen, M. Thulin, M. Thurston, B. Thuy, A. Todaro, M. Todorov, R. Toonen, B. M. Torke, S.-Y. Tsai, M. Turiault, J. R. G. Turner, T. Turner, X. Turon, S. Tyler, UCD Community, UNITE Community, D. A. Uchima Taborda, P. Uetz, J. M. Ulmer, J. S. Usma Oviedo, J. Utria Suarez, J. Vacelet, D. Vachard, W. Vader, G. Valls Domedel, X. Van der Burgt, L. Vandepitte, B. Vanhoorne, V. Vargas-Canales, M. Vatanparast, D. Vaulot, T. Verhoeff, R. Verovnik, P. Vieira, R. Vila, F. A. Villa-Navarro, M. Vilà, A. Vliegenthart, K. Vongphayloth, R. Vonk, R. Väinölä, T. WANG, N. Wahlberg, G. Walker-Smith, T. C. Walter, N. Wambiji, D. Wanke, M. Warren, L. Watling, H. Weaver, J. Webb, W. C. Welbourn, T. Wesener, C. Whipps, K. White, M. Wiemers, N. Wilding, G. Williams, A. J. G. Wilson, D. E. Wilson, P. Wing, S. Winitsky, M. Winter, C. C. Wirth, M. Wojciechowski, S. Woodman, J. Xavier, T. Yi, M. Yoder, D. S. K. Yu, N. Yunakov, P. Yésou, M. ZHAO, J. Zahniser, A. Zaiko, F. Zapata, L. A. Zapata, L. A. Zapata Padilla, S. Zaragoza-Caballero, W. Zeidler, R. Zhang, F. Zinetti, J.-L. d'Hondt, G. J. de Moraes, A. B. R. de Oliveira, N. de Voogd, J. del Campo, M. G. del Río, iNaturalist contributors, T. van Haaren, E. J. van Nieukerken, L. van Ofwegen, R. van Soest, C. A. M. van Swaay, S. van der Meij, O. Şentürk, American Society of Mammalogists, Cornell Lab of Ornithology, Cornell University, ITIS, International Barcode of Life project (iBOL), International Committee on Taxonomy of Viruses (ICTV), Legume Phylogeny Working Group (LPWG), Natural History Museum Bern, Paleobiology Database contributors, Smithsonian Institution National Museum of Natural History, USDA, Agricultural Research Service, National Germplasm Resources Laboratory, WoRMS Editorial Board, World Flora Online, Catalogue of Life (2026). https://doi.org/10.48580/dgxsq.
